# High Throughput Designing and Mutational Mapping of RBD-ACE2 Interface Guide Non-Conventional Therapeutic Strategies for COVID-19

**DOI:** 10.1101/2020.05.19.104042

**Authors:** Aditya K. Padhi, Parismita Kalita, Kam Y. J. Zhang, Timir Tripathi

## Abstract

Considering the current status of the SARS-CoV-2 pandemic, sequence variations and possibly structural changes in the rapidly evolving SARS-CoV-2 is highly expected in the coming months. The SARS-CoV-2 spike (S) protein is responsible for mediating viral attachment and fusion with cell membranes. Mutations in the receptor-binding domain (RBD) of the S-protein occur at the most variable part of the SARS-CoV-2 genome, and specific sites of S-protein have undergone positive selection impacting the viral pathogenicity. In the present work, we used high-throughput computation to design 100,000 mutants in RBD interfacial residues and identify novel affinity-enhancing and affinity-weakening mutations. Our data suggest that SARS-CoV-2 can establish a higher rate of infectivity and pathogenesis when it acquires combinatorial mutations at the interfacial residues in RBD. Mapping of the mutational landscape of the interaction site suggests that a few of these residues are the hot-spot residues with a very high tendency to undergo positive selection. Knowledge of the affinity-enhancing mutations may guide the identification of potential cold-spots for this mutation as targets for developing a possible therapeutic strategy instead of hot-spots, and vice versa. Understanding of the molecular interactions between the virus and host protein presents a detailed systems view of viral infection mechanisms. The applications of the present research can be explored in multiple antiviral strategies, including monoclonal antibody therapy, vaccine design, and importantly in understanding the clinical pathogenesis of the virus itself. Our work presents research directions for the exploitation of non-conventional solutions for COVID-19.

## 1. INTRODUCTION

As of May 16, 2020, the novel severe acute respiratory syndrome coronavirus-2 (SARS-CoV-2), the causal agent of the coronavirus disease (COVID-19), has infected more than 4.6 million individuals and has claimed the lives of more than 300,000 people across 212 countries globally. The virus has a positive, single-stranded genome that is over 29 kb in length [1]. The spike (S) protein, being the sole protein responsible for receptor binding and viral entry, stands as the primary target of neutralizing antibodies. The S-protein is a class I fusion protein, with each S protomer consisting of S1 and S2 domains. The receptor-binding domain (RBD) of SARS-CoV-2 (SARS-CoV-2-RBD) is located at the C-terminus of the S1 domain of S-protein. Cleavage of the S-protein by host proteases mediates subsequent virus-cell fusion and initiates infection. Studies show that the SARS-CoV-2-RBD binds to the human angiotensin-converting enzyme-2 (ACE2) receptor with a higher affinity (~10 to 15 nM), justifying the efficient transmission of SARS-CoV-2 [2, 3]. A cryo-EM trimeric structure of the SARS-CoV-2 S-protein in its prefusion conformation has been determined recently [2]. The prefusion trimer is destabilized upon ACE2 binding leading to the formation of a stable post-fusion state. Several structures of the SARS-CoV-2-RBD-ACE2 complex have also been elucidated, and the interfacial residues have been identified [4–6]. The SARS-CoV-2-RBD consists of seven β sheets (β1-β7) with short connecting helices and loops forming the core structure. The β5 and β6 strands, α4 and α5 helices and loops are present as a short extended insertion between the β4 and β7 strands. This extended insertion is the receptor-binding motif that contains most interfacial residues. The residual interactions involved in RBD-ACE2 complex formation are also identified and rationalized, highlighting the crucial differences with other SARS-CoVs. Thus, S-protein represents the crucial therapeutic target, and focusing on the critical understanding of the molecular mechanism of RBD-ACE2 interaction and finding potential weaknesses can be exploited for therapeutic purposes.

Several studies have identified specific replacement of critical amino acids between the S-protein of various SARS-CoVs that have rendered the human cells susceptible to the SARS-CoV and SARS-CoV-2 infection [2, 5–8]. The amino acid sequence variations between the RBD of the SARS-CoV-2 and other SARS-CoVs have been elucidated [1, 9]. The RBD of SARS-CoV-2 and SARS-CoV possess a similar structure, despite amino acid variations at key residues [1]. However, mutations in the RBD occur at the most variable part of the SARS-CoV-2 genome. Evolutionary genomics data suggest that the SARS-CoV-2 S-protein has gained several mutations that increased its binding affinity towards ACE2 by ~10-15-fold compared to the S-protein of SARS-CoV [9, 10]. The affinity of RBD for its receptor ACE2 is one of the major determinants of the virus transmission rate. While SARS-CoV-2 has a relatively low mutation rate, genome data show the emergence of novel mutations with altered virulence. SARS-CoV-2 with D614G mutation is presently the most common mutant form of the virus, accelerating the spread of the pandemic in several countries [11, 12]. A molecular dynamics simulation study on the effect of RBD mutations, from recently reported SARS-CoV-2 genomes, showed that V367F, W436R, and D364Y displayed a higher affinity towards the ACE2 receptor [13]. On the contrary, in another study, the conserved residue R408 of RBD of S-protein present in the interface of RBD and ACE2 receptor, upon mutation to isoleucine was found to show reduced binding affinity by disrupting a hydrogen bond formed with glycan positioned at 90N from ACE2 [14]. Such findings highlight the importance of monitoring the mutational dynamics of the SARS-CoV-2 genome. Accumulation of more genome sequence data shows the prospects of identification of several such mutants that could explain the selective pressure on SARS-CoV-2 evolution.

Since the interactions between RBD and ACE2 are critical for viral entry and pathogenesis, it is vital to identify critical interfacial residues that, upon mutation, either protect or render individuals more susceptible to the virus. Two recent studies identified such ACE2 variants and reported mutations that enhanced or decreased binding to S-protein, thereby potentially altering RBD-ACE2 interactions [15, 16]. However, the applications of their work are limited as natural variations in ACE2 in human populations are extremely rare. As a more rational approach to the problem, we decided to identify combinatorial mutations in the RBD interfacial residues that either increase or decrease its affinity towards the ACE2 receptor. This is crucial as the rapid spread of the virus provides it an ample opportunity for natural selection to act upon rare but favorable mutations. In the present work, we used high-throughput computational approaches to characterize the detailed RBD and ACE2 interaction networks and identify novel affinity-enhancing and affinity-weakening mutations. The data provide crucial insights into interaction and mutational dynamics between RBD and ACE2 that could be explored for developing strategies for novel and effective therapeutics.

## 2. METHODS

### 2.1. Identification of interacting residues at the interface of RBD-ACE2 complex

The crystal structure of the RBD-ACE2 complex (PDB ID: 6LZG) was retrieved from RCSB Protein Data Bank and used for obtaining the critical interacting residues of the RBD with ACE2 [3]. A total of 24 unique residues of RBD were identified to be interacting with several residues of ACE2 (Table 1, Figure 1). Subsequently, the mutation sensitivity profile of RBD was obtained using MAESTROweb to determine positions where mutations are predicted to have little effect on the stability, or if they are predominantly stabilizing or destabilizing [17].

**Table 1.** Interacting residues between ACE2 and RBD obtained from the crystal structure of RBD-ACE2 complex.

| **ACE2** | **RBD** | **ACE2** | **RBD** | **ACE2** | **RBD** | **ACE2** | **RBD** | **ACE2** | **RBD** | **ACE2** | **RBD** | **ACE2** | **RBD** |
| --- | --- | --- | --- | --- | --- | --- | --- | --- | --- | --- | --- | --- | --- |
| THR27 | TYR489 | GLN42 | GLN498 | ARG393 | TYR505 | LYS31 | PHE490 | GLN24 | GLY476 | GLU37 | TYR505 | GLN42 | TYR449 |
| GLN24 | ASN487 | ASP355 | ASN501 | HIS34 | TYR453 | LEU79 | GLY485 | TYR41 | ASN501 | LEU45 | THR500 | HIS34 | ARG403 |
| TYR83 | ASN487 | THR324 | GLY502 | THR27 | TYR473 | THR324 | VAL503 | ASP355 | GLY502 | GLN24 | TYR489 | GLN42 | GLY446 |
| GLY354 | TYR505 | HIS34 | GLN493 | GLY354 | VAL503 | GLY354 | ASN501 | ASP355 | THR500 | LEU79 | PHE486 | TYR41 | GLN498 |
| LYS31 | TYR489 | TYR83 | PHE486 | ASP38 | TYR495 | ASP38 | GLN498 | ARG357 | THR500 | PHE28 | TYR489 | GLY354 | GLY502 |
| ALA386 | TYR505 | ASP38 | TYR449 | LYS31 | PHE456 | ASP30 | PHE456 | ASP30 | LEU455 | GLN24 | SER477 | THR27 | PHE456 |
| LYS353 | ASN501 | ASN330 | THR500 | LYS353 | GLY496 | TYR41 | THR500 | LYS353 | TYR495 | LYS31 | LEU455 | GLN24 | ALA475 |
| LYS353 | GLN498 | ASP38 | GLY496 | MET82 | PHE486 |  |  |  |  |  |  |  |  |

**Figure 1.**
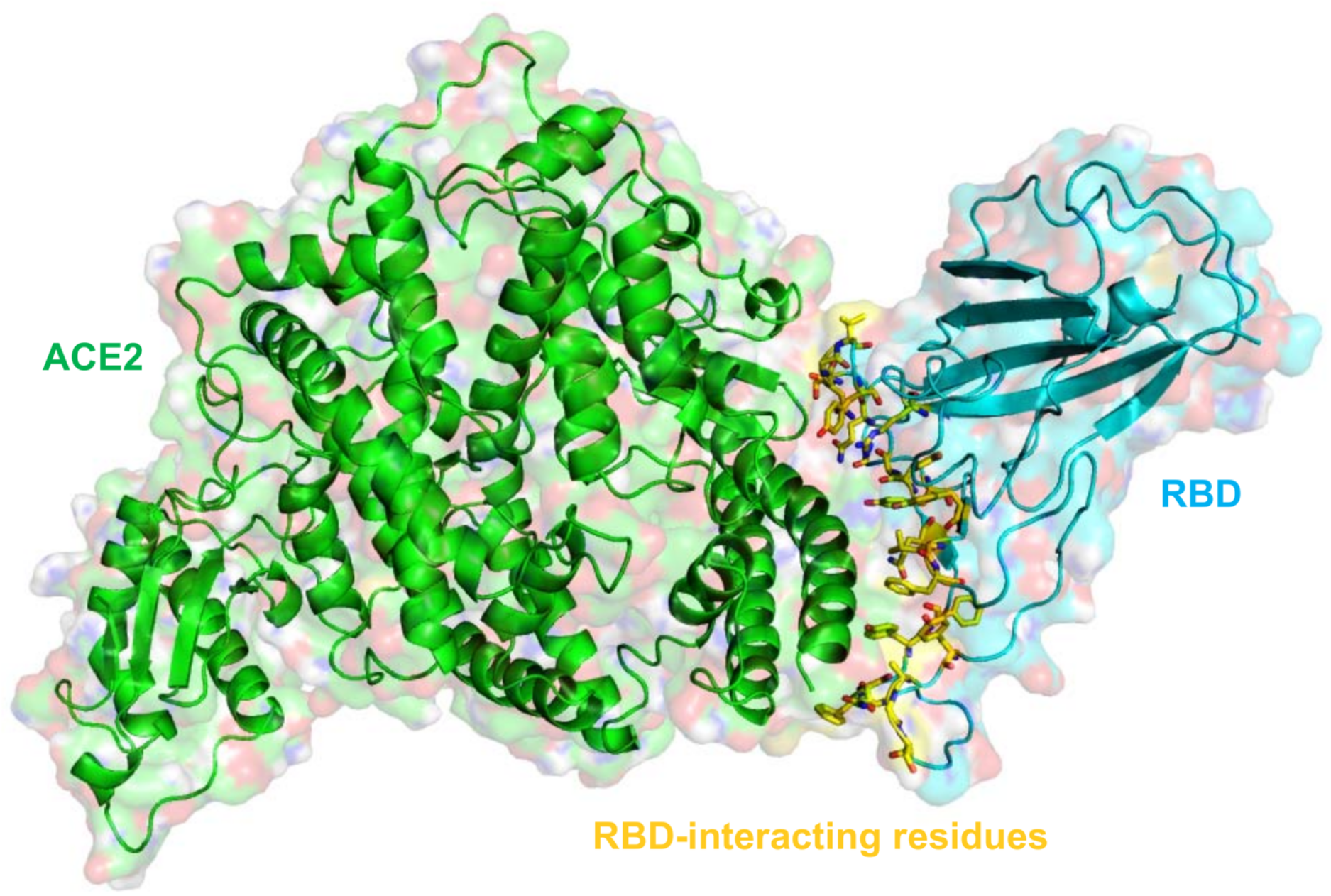
Crystal structure of ACE2-RBD complex showing interacting residues. The crystal structure of ACE2 in complex with RBD of SARS-CoV-2 is shown along with the interacting residues of RBD. The ACE2 and RBD are shown in cartoon and surface representations in green and cyan color, respectively, and the interacting residues of RBD are shown in yellow sticks.

### 2.2. Structure refinement for Rosetta interface design

The resulting structure of the RBD-ACE2 complex was optimized using Schrödinger Maestro (Schrödinger Release 2016–4: Maestro, Schrödinger, New York) and used for structural preparation and refinement by Rosetta relax, the main protocol for all-atom refinement of structures in the Rosetta force-field [18]. In this step, ten models were generated, and the fullatom relaxed structure with the lowest energy was used for subsequent interface design experiments.

### 2.3. Flexible backbone interface design and binding affinity calculation

Rosetta macromolecular modeling suite was used to carry out the interface design experiments [19]. The RBD of the RBD-ACE2 complex was redesigned, keeping the backbone flexible and using Rosetta’s full-atom scoring function. In this step, only the 24 residues of RBD known to interact with ACE2 were redesigned with eighteen other amino acids (except cysteine to prevent disulfide bond formation during interface design). During this stage, the remaining residues of RBD and ACE2 were allowed to repack without design. A total of 100,000 designs were generated where ten cycles were carried out to generate a structure. In the design simulations, a modified RosettaScripts was used to design the interface residues that consider the Backrub flexible backbone modelling method [20]. The designs were then sampled by Monte-Carlo simulated annealing using the Rosetta all-atom force field. For a detailed structural and quantitative analysis with a focus on in cerebro examination of the resulting designs, the Rosetta total score, root mean square deviation (RMSD) from the starting structure, Rosetta DDG representing the binding affinity between designed RBD with ACE2, and the percentage sequence identity from the starting sequence were obtained.

### 2.4. Validating the interface design protocol on high-frequency RBD mutations

To test the accuracy and reliability of our Rosetta interface design protocol, we took two mutants of RBD (G476S and V483A) into consideration, which were high-frequency RBD mutations identified from a study involving 1609 different strains of the virus[14, 21]. In this stage, we designed Gly476 and Val483 residues, whereas the remaining residues were allowed to repack without design. A total of 10,000 designs per designed position were generated and were analyzed to verify the scoring and ranking of the mutants.

### 2.5. Sequence logos of top-ranked affinity-enhancing and affinity-weakening designs

The frequency and types of designed residues in the top-ranked affinity-enhancing and affinity-weakening designs were obtained and plotted using WebLogo that gives a graphical representation of an amino acid multiple sequence alignment [22]. In each plot, X-axis denotes the residue index, and the overall height of the stack in the Y-axis denotes the sequence conservation at that position. The height of symbols within the stack indicates the relative frequency of each amino acid at that position.

### 2.6. PRODIGY-based binding affinity calculation

The Rosetta based interface designs were then subjected to the calculation of the binding affinities between RBD and ACE2 using PROtein binDIng enerGY prediction (PRODIGY), a predictor of binding affinity in a protein-protein complex from their 3D structures [23]. In this method, after the 3D model is supplied with a suitable temperature (default of 25°C), the binding affinity is computed. From the Rosetta designed structures, the binding affinity (ΔG) and the number of intermolecular contacts were obtained for analysis. Detailed decomposition of intermolecular contacts was then obtained for the top five affinity-enhancing and affinity-weakening designs.

### 2.7. Calculation of intermolecular interactions governing the affinity-enhancing and affinity-weakening mutants

The intermolecular interactions between ACE2 and RBD for the top-ranked affinity-enhancing and affinity-weakening designs were obtained using Arpeggio web-server [24]. The total number of interactions obtained represents the sum of the number of van der Waals, proximal, polar contacts, hydrogen bonds, aromatic contacts, hydrophobic contacts, and carbonyl interactions. As a final step, residue level interactions between ACE2 and RBD interface for the top-ranked affinity-enhancing and affinity-weakening designs obtained by the DIMPLOT program of LigPlot+v1.4.5 and were analyzed in Pymol.

## 3. RESULTS

### 3.1. Interacting residues of RBD-ACE2 complex and mutation sensitivity profile

The crystal structure of the RBD-ACE2 complex revealed that the interface between them is stabilized by 59 interactions in total (Table 1). Among the 59 interactions, 23 residues from RBD establish intermolecular contacts with the ACE2 interface (Figure 1). We were curious to learn how the residues of RBD affect the stability when they are mutated systematically to other residues. To understand this, we obtained a mutation sensitivity profile, where the impact of each mutation at each possible position is visualized. It was found that the interface residues of RBD that interact with ACE2 are not highly stable at many positions with ΔΔG_pred_>0 (Supplementary Figure S1). Some of the hot-spot residues such as R403, G446, Y453, L455, F456, Y473, G485, Y489, F490, Y495, N501 and G502 with ΔΔG_pred_>0 suggests they are relatively unstable and likely more prone to mutation. It is noteworthy that some of these native RBD residues are already found to be mutated in the SARS-CoV-2 genome, as reported in the CoV-GLUE database, which publishes information on amino acid replacements, insertions, and deletions from genome sequences sampled from the pandemic (http://cov-glue.cvr.gla.ac.uk/#/replacement). For instance, G446 to S, Y453 to S, F456 to L, Y473 to D/H, Y489 to S, F490 to L/S, Y495 to S/N, N501 to T, etc. This further confirms our mutational sensitivity profiling. However, this does not consider the flexibility of the backbone residues while suggesting the sensitivity profile, owing to which, we opted for interface designing focused on the RBD of SARS-CoV-2 to gain further structural insights.

### 3.2. Rosetta flexible backbone interface design and various structural features

To gain insights on how the RBD of SARS-CoV-2 can modulate the interaction with ACE2, we designed the interface of RBD-ACE2 with a focus on the RBD specifically. In our design experiments, we allowed the backbone flexibility of RBD during design, where 24 interacting residues of RBD were mutated to other remaining amino acids except cysteine. The 100,000 generated designs were first sampled by calculating the total scores versus RMSD of the designs. We found that >50% of the topmost total-scored designs exhibited RMSD <2Å, suggesting even after mutations of the interface residues of the RBD, the whole complex doesn’t deviate much from the crystal structure, thus maintaining structural integrity (Figure 2A).

**Figure 2.**
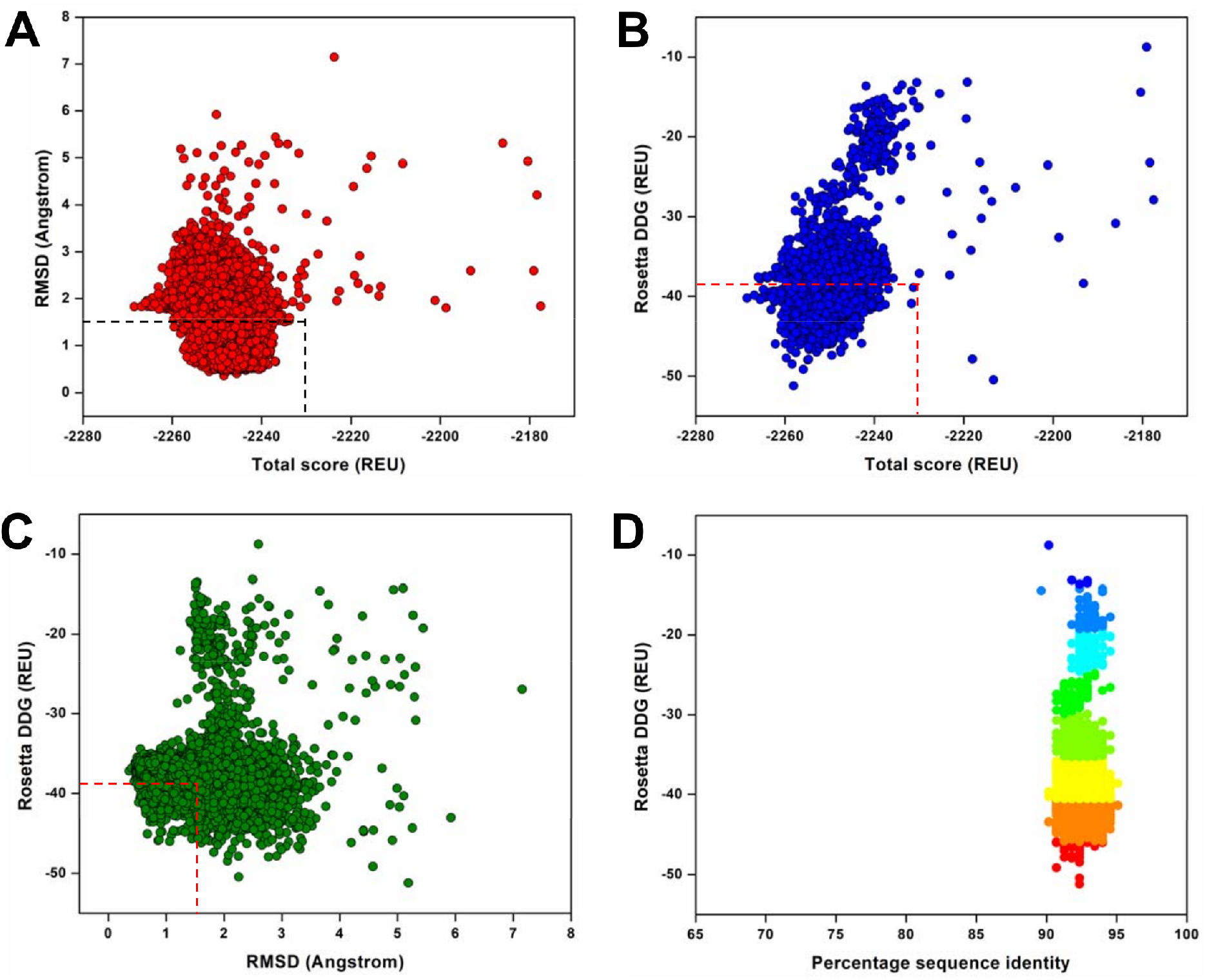
Structural and quantitative parameters from Rosetta interface design experiments. (A) Rosetta total score (Rosetta Energy Units, REU) versus RMSD (Å) of the 100,000 design structures of ACE2-RBD complexes obtained from flexible backbone design, (B) Rosetta total score (REU) versus Rosetta DDG (REU) representing the binding affinities between RBD-ACE2 for the computed designs, (C) Rosetta DDG (REU) versus RMSD (Å) of the designs, where the dotted boxes denote the control values where the RBD interacting residues were only repacked and not designed, (D) Rosetta DDG (REU) versus percentage sequence identity of the generated designs against the native RBD sequence.

Secondly, we calculated the Rosetta DDG of the designs and plotted them against the total scores. Importantly, we also observed that the designs with top-scored DDG/binding affinities have higher total scores, indicating that the designs with mutations at the interface interacting residues of RBD converge well during the simulations (Figure 2B, Supplementary Figure S2). For distinguishing the designs among themselves, we took the ACE2-RBD complex and only repacked (no designing) the same set of interacting residues of RBD and compared it with the DDG and total score as control (Figure 2B). From this analysis, we further confirmed that many top-scored designs exhibit much higher binding affinities than the control in our design experiments.

Thirdly, our computation of Rosetta DDG versus the RMSD further demonstrated the convergence of the design simulations and better affinity variants of RBD (Figure 2C). As expected, here we found that some of the better binding affinity designs (out of the red box) exhibit relatively higher RMSDs (from 1.5-3.5 Å) but still retain higher binding affinities, suggesting that certain mutations have changed the backbone conformation resulting in slightly increased deviation in RMSDs (Figure 2C).

Our final evaluation of binding affinities in comparison to percentage sequence identity revealed that the designs exhibiting higher binding affinities most often contain sequences with 90-95% identity to that of native sequence (Figure 2D). However, the RBD designs having lower binding affinities showed relatively low percent sequence identity, suggesting their mutational profiles are less diverse, thus ensuing low stability as a whole when the RMSDs and total scores are taken into consideration (Figure 2D). This also suggests that SARS-CoV-2 can establish a higher rate of infectivity and pathogenesis when it acquires combinatorial mutations (not just single point mutations) at the RBD of the RBD-ACE2 complex during evolutionary pressure.

### 3.3. Validation of Rosetta flexible backbone interface design

To validate our interface design methodology, two residues of RBD, such as G467 and V483, were designed and evaluated to see whether our methodology is able to score and rank G476S and V483A at the top because they have the highest occurrence in several isolates across diverse populations. Our analysis of binding affinities versus total score revealed that the G467S mutant scores better in both binding affinity and total score as compared to V483A mutant in the design experiments (Supplementary Figure S3). However, both the designs scored much better in the design simulations when compared to other residues that are sampled during the designs and rank-ordered. This suggests that our design protocol and workflow are capable enough to score, rank order, and distinguish the favorable and unfavorable RBD mutants for binding with ACE2 (Supplementary Figure S3).

### 3.4. Diversity of top-ranked affinity-enhancing and affinity-weakening designs

We subsequently divided the designs into two groups, such as the affinity-enhancing designs and the affinity-weakening designs. The affinity-enhancing designs had higher total scores in addition to better binding affinity. The analysis of 50 top-scored RBD designs from each group revealed a landscape of mutations at the interacting residues of RBD (Figure 3). It was observed that while certain positions of the native RBD such as Arg403, Gly446, Gly476, Gly485, Phe490, Tyr495, Gly496, and Gly502 are sampled to almost identical residues in both the groups, they varied most at other positions of the remaining residues. However, residues Phe456, Tyr473, Phe486, Tyr489, Gln498, Thr500, and Asn501 were mutated to unique residues in both groups, which drive the affinity enhancing or weakening characteristics towards ACE2 (Figure 3). Some of these residues were found to be the hot-spot residues in mutation sensitivity profile (Supplementary Figure S1), altogether suggesting these are the ones probably to be used by the SARS-CoV-2 for its infection, propagation, and survival from its evolutionary and immunological pressure.

**Figure 3.**
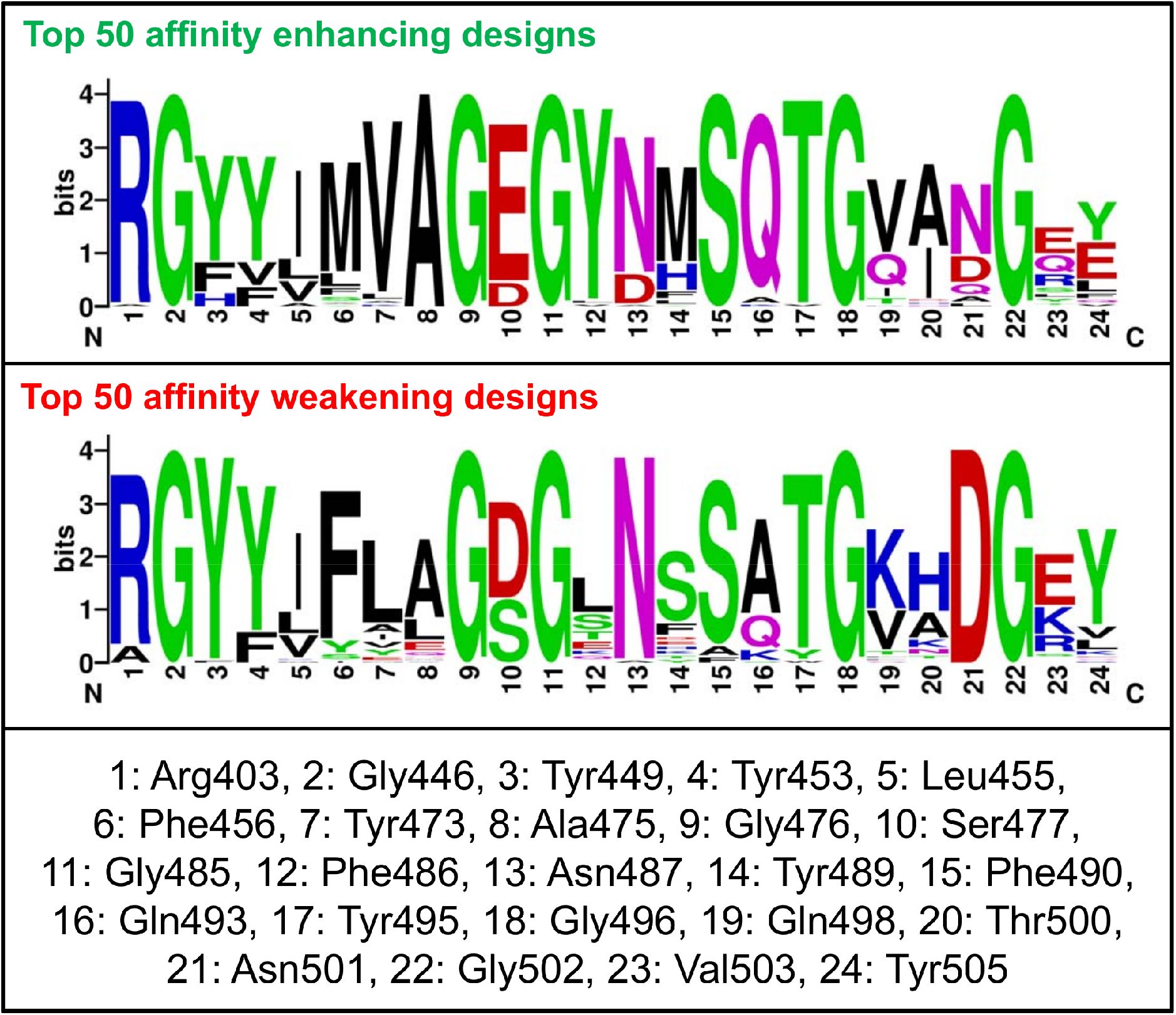
Sequence logos showing the frequency of designed RBS residues. Sequence logos of top 50 affinity-enhancing and affinity-weakening designs generated from the Rosetta interface flexible design approach for the RBD of SARS-CoV-2. The integers on the X-axis representing the corresponding native residues and their identities are shown in the bottom panel. The ‘bits’ represent the overall height of the stack in the Y-axis with the sequence conservation at that position.

### 3.5. RBD designs-ACE2 binding affinity calculation using PRODIGY

Next, we computed the binding affinities of the RBD designs with ACE2 obtained from Rosetta simulations. We found that the binding affinities ranged from −8.8 kcal/mol to −12.5 kcal/mol across the designs (Figure 4). When the binding affinities were compared, many of the designs scored higher in both methodologies, suggesting that they formed a higher number of interactions and contacts with ACE2 when certain positions of the RBD were mutated. An analysis of the number of intermolecular contacts versus the binding affinities (Figure 4), further confirmed the assumption that Rosetta generated top-scored higher binding affinity designs can facilitate SARS-CoV-2 to establish stronger interactions with ACE2. Detailed decomposition of intermolecular contacts of the top five affinity-enhancing and affinity-weakening designs further demonstrated that the numbers of intermolecular contacts, charged-apolar contacts, polar-polar contacts, apolar-polar contacts, and apolar-apolar contacts primarily drive the affinity-enhancing or affinity-weakening RBD mutants towards ACE2 (Tables 2 and 3).

**Figure 4.**
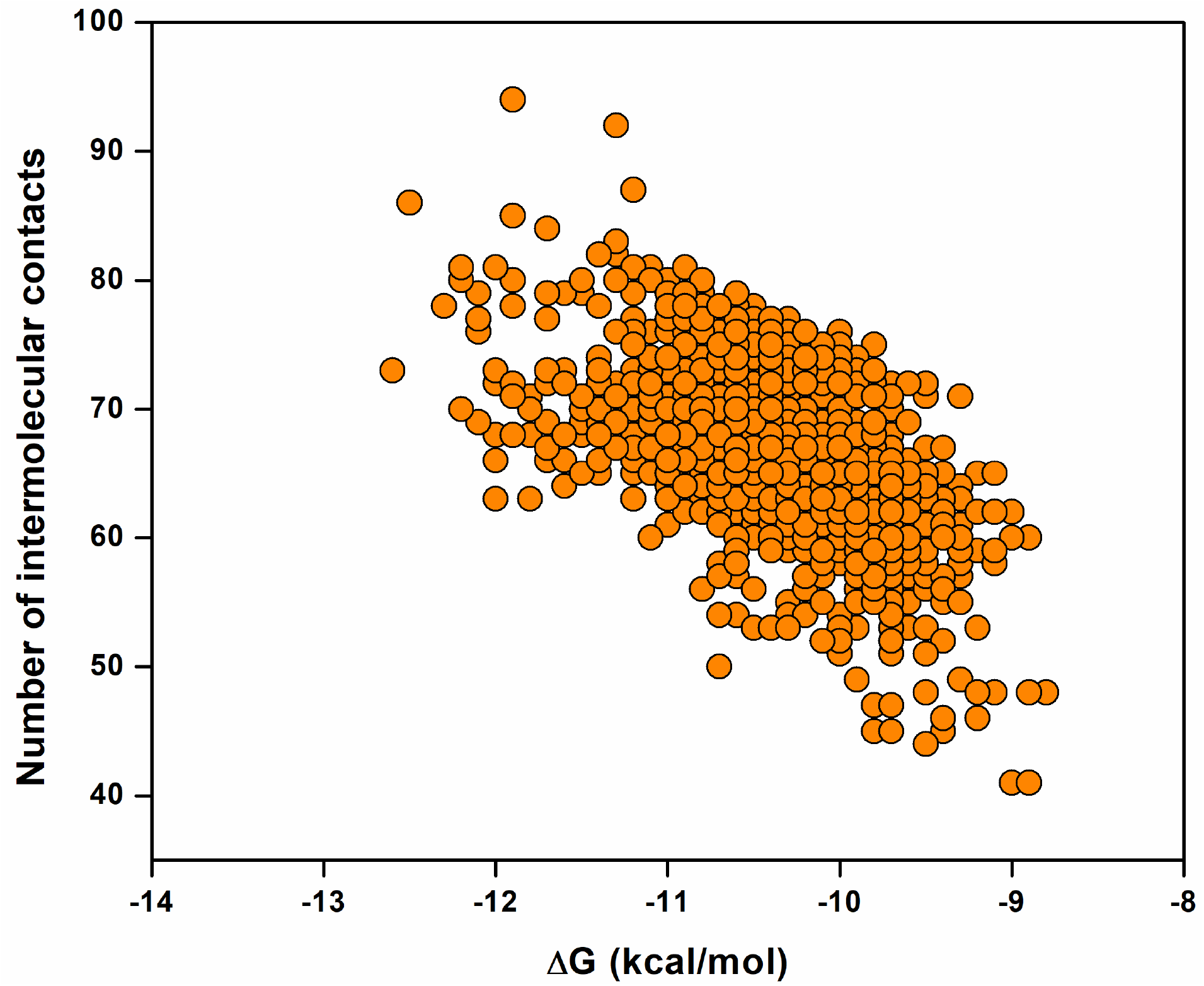
PRODIGY derived binding affinities and intermolecular contacts. The binding affinities of the designs represented in (ΔG) versus the number of intermolecular contacts formed with ACE2 are shown.

**Table 2.** Detailed decomposition of the intermolecular contacts of top five affinity-enhancing designs.

| **RBD-**<br>**ACE2**<br>**interface**<br>**designs** | **No. of**<br>**intermolecular**<br>**contacts** | **No. of**<br>**charged-**<br>**charged**<br>**contacts** | **No. of**<br>**charged-**<br>**polar**<br>**contacts** | **No. of**<br>**charged-**<br>**apolar**<br>**contacts** | **No. of**<br>**polar-**<br>**polar**<br>**contacts** | **No. of**<br>**apolar-**<br>**polar**<br>**contacts** | **No. of**<br>**apolar-**<br>**apolar**<br>**contacts** | **% of**<br>**apolar**<br>**NIS**<br>**residues** | **% of**<br>**charged**<br>**NIS**<br>**residues** | **Predicted**<br>**binding**<br>**affinity**<br>**(kcal/mol)** |
| --- | --- | --- | --- | --- | --- | --- | --- | --- | --- | --- |
| Design-1 | 86 | 8 | 11 | 25 | 3 | 19 | 20 | 34.73 | 28.14 | -12.5 |
| Design-2 | 80 | 6 | 9 | 22 | 4 | 20 | 19 | 34.29 | 27.97 | -12.2 |
| Design-3 | 81 | 4 | 9 | 27 | 3 | 18 | 20 | 34.38 | 28.08 | -12.2 |
| Design-4 | 79 | 6 | 10 | 26 | 3 | 17 | 17 | 34.38 | 28.23 | -12.1 |
| Design-5 | 77 | 5 | 13 | 23 | 3 | 19 | 14 | 34.59 | 28.27 | -12.1 |

**Table 3.** Detailed decomposition of the intermolecular contacts of top five affinity-weakening designs.

| **RBD-**<br>**ACE2**<br>**interface**<br>**designs** | **No. of**<br>**intermolecular**<br>**contacts** | **No. of**<br>**charged-**<br>**charged**<br>**contacts** | **No. of**<br>**charged-**<br>**polar**<br>**contacts** | **No. of**<br>**charged-**<br>**apolar**<br>**contacts** | **No. of**<br>**polar-**<br>**polar**<br>**contacts** | **No. of**<br>**apolar-**<br>**polar**<br>**contacts** | **No. of**<br>**apolar-**<br>**apolar**<br>**contacts** | **% of**<br>**apolar**<br>**NIS**<br>**residues** | **% of**<br>**charged**<br>**NIS**<br>**residues** | **Predicted**<br>**binding**<br>**affinity**<br>**(kcal/mol)** |
| --- | --- | --- | --- | --- | --- | --- | --- | --- | --- | --- |
| Design-1 | 48 | 9 | 13 | 10 | 0 | 6 | 10 | 34.38 | 28.42 | -8.8 |
| Design-2 | 48 | 13 | 6 | 14 | 1 | 4 | 10 | 34.62 | 28.11 | -8.9 |
| Design-4 | 41 | 7 | 6 | 14 | 0 | 5 | 9 | 34.57 | 27.74 | -8.9 |
| Design-5 | 48 | 13 | 6 | 14 | 1 | 4 | 10 | 34.62 | 28.11 | -8.9 |
| Design-3 | 41 | 11 | 7 | 12 | 1 | 6 | 4 | 34.90 | 27.54 | -9.0 |

### 3.6. Visualization and computation of intermolecular interactions for the affinity-enhancing and affinity-weakening mutants

In the end, we were interested in visualizing and computing the types of intermolecular interactions that play a role in inducing the improved or reduced affinity of the RBD designs towards ACE2 (Figure 5). For this, we took two designs representing the top-ranked affinity-enhancing and affinity-weakening characteristics. In cerebro analysis revealed a significantly higher number of interactions with a much compact binding interface for the affinity-enhancing Design-1 (Figure 5A) as compared to affinity-weakening Design-1 (Figure 5B). The number and types of intermolecular interactions between both the designs demonstrate this feature (Table 4). When comparing how the intermolecular interactions of the RBD-ACE2 crystal structure contrast with our affinity-enhancing Design-1, we found that although the crystal structure has a higher number of interactions, the total number of intermolecular interactions in the designed affinity-enhancing mutants surpassed the crystal structure, mainly due to the increased number of van der Waals and proximal interactions (Table 4). Finally, the residue level interactions between ACE2 and RBD for the top-ranked affinity-enhancing and affinity-weakening designs were obtained by the DIMPLOT program, as shown in Supplementary Figure S4.

**Figure 5.**
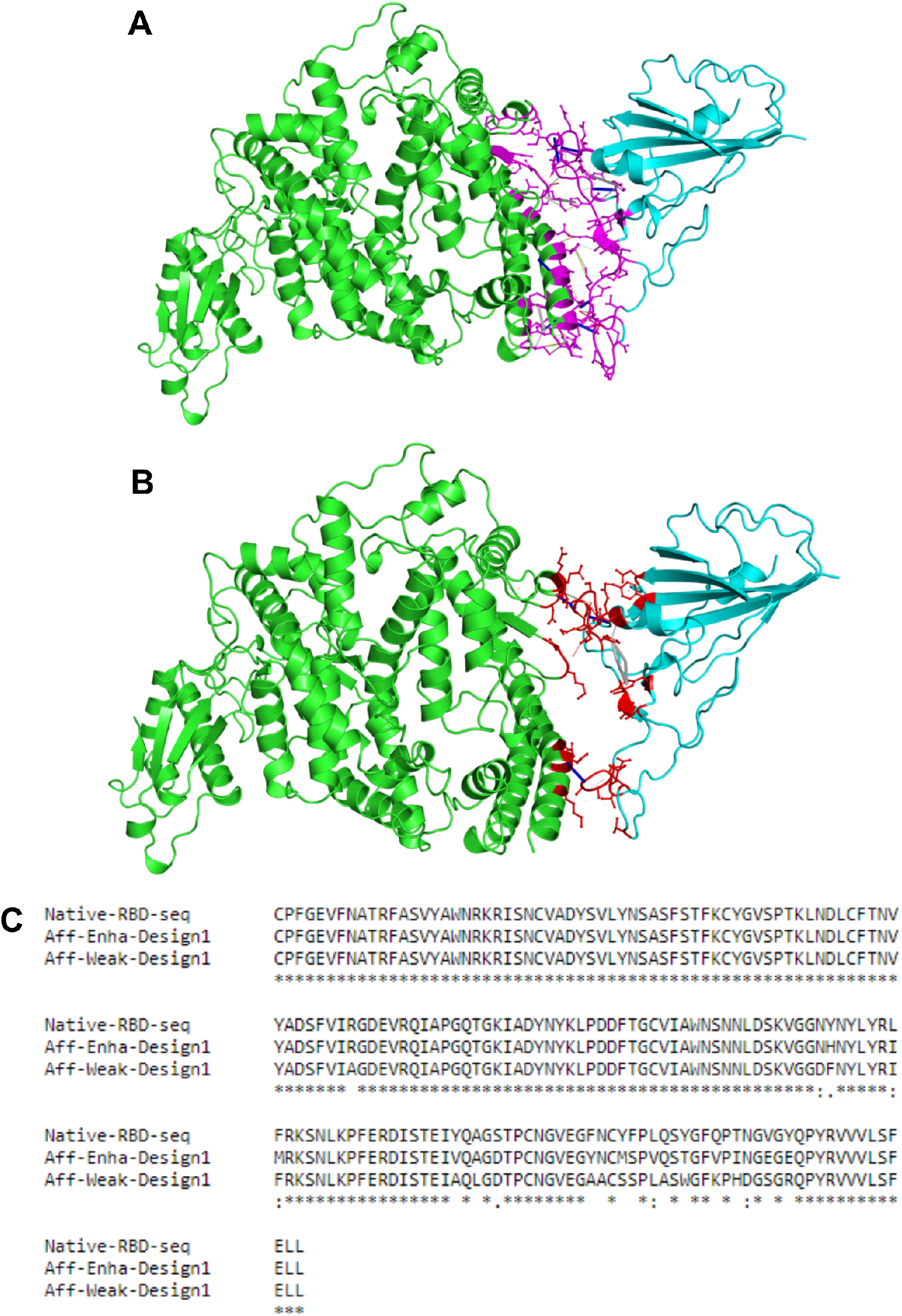
Key interactions obtained from the top affinity-enhancing and affinity-weakening designs. Key intermolecular interactions between ACE2-RBD for the (A) top-ranked affinity-enhancing design-1 and (B) top-ranked affinity-weakening design-1 are shown. In both (A) and (B), ACE2 is shown in green color, and RBD is shown in cyan color. In panel A, the interacting residues and binding site are shown in purple stick lines, and in panel B, the interacting residues and binding site are shown in red stick lines. Various types of intermolecular interactions such as van der Waals interactions, proximal interactions, polar contacts, hydrogen bonds, aromatic contacts, hydrophobic contacts, carbonyl interactions, and amide-amide interactions are shown in yellow, grey, red, white dashed, white long-dashed, green dashed, black-white dashed and in blue dashed lines respectively. (C) Multiple sequence alignment of native RBD sequence along with the affinity-enhancing design-1 and affinity-weakening design-1 showing the differences in mutated residues.

**Table 4.** Detailed intermolecular interactions formed between RBD and ACE2 for the top affinity-enhancing and affinity-weakening designs compared to the crystal structure.

| **Types of interactions** | **Design-1 (Affinity-enhancing)** | **Design-1 (Affinity-weakening)** | **RBD-ACE2 crystal structure** |
| --- | --- | --- | --- |
| Van Der Waals interactions | 10 | 1 | 8 |
| Proximal interactions | 829 | 345 | 763 |
| Polar contacts | 20 | 5 | 19 |
| Hydrogen bonds | 15 | 4 | 15 |
| Aromatic contacts | 2 | 0 | 9 |
| Hydrophobic contacts | 44 | 5 | 61 |
| Carbonyl interactions | 1 | 1 | 1 |
| Total number of interactions | 921 | 361 | 876 |

## 4. DISCUSSION

In the last two decades, three major outbreaks are caused by a novel human CoVs-SARS-CoV in 2002, Middle East respiratory syndrome CoV (MERS-CoV) in 2012, and now in 2019 SARS-CoV-2. SARS-CoV-2 is one of the most transmissible CoVs identified to date. The SARS-CoV-2 S-protein binds specifically to the transmembrane ACE2 receptor to gain entry into host cells. Thus, the blocking of S-protein to ACE2 receptor interaction, and cell fusion is one of the key strategies for antiviral development. The binding free energy of RBD to ACE2 is the key determinant of SARS-CoV-2 transmissibility. The higher affinity of SARS-CoV-2-RBD to ACE2 is attributed to the presence of an extended network of favorable hydrogen bond networks between the interacting partners. Interestingly, the SARS-CoV-2-RBD displays a 10-20 fold stronger affinity for ACE2 compared with the RBD of other SARS-CoVs [2, 3]. This suggests that the SARS-CoV-2 encodes distinct epitope properties in the RBD from other SARS-CoVs.

The RNA viruses evolved a specific strategy to interact with the host membrane. They use protein-binding interaction motifs specific to RNA viruses [25]. Multiple virus and host-dependent processes contribute to viral diversity by encouraging mutations in the viral proteins to adapt to a new host and environment [26]. Mutations in the viral binding proteins modulate infectivity and alter antigenicity leading to species specificity, as in the case of SARS-CoV-2-RBD. The spike gene of the SARS-CoV-2 may be recombined with the other SARS-CoVs due to convergent evolution (with its relatively high mutation rate), which is the very nature of SARS-CoVs [27]. Under natural selection and evolutionary pressure, the SARS-CoV-2-RBD has undergone several mutations conferring stability to the viral particle [1, 9, 28]. Pieces of evidence are accumulating on viruses undergoing positive selection suggesting the emergence of a more transmissible form of SARS-CoV-2 [29–31]. However, the mutation rate of SARS-CoV-2 is not particularly high with respect to other SARS-CoVs, but they do accumulate with time. Thus, as the epidemic progresses, accumulations of further mutations due to positive selective pressure could favor an enhancement of pathogenicity and transmission of SARS-CoV-2. This may accelerate the risk for antigenic drift and the accumulation of immunologically significant mutations in the population.

To understand the complex mechanisms of the infection process, computational analysis of underlying protein-protein interaction (PPI) networks may provide crucial insights for designing novel and effective therapeutics. To modulate SARS-CoV-2-RBD interactions with ACE2, we applied high-throughput mutational screening. We observed that in several mutants, the binding affinities had similar binding interface sizes but altered RBD-ACE2 interactions. Determinants of binding affinity include ‘hot spots’, i.e., residues at critical positions in the binding interface for which mutations cause a significant reduction in affinity. Mutations at other positions in the PPI interfaces are known as ‘cold spots’ and do not significantly affect or may sometime increase the binding affinity [32]. Knowledge of the direction of mutational pressure may help design more effective therapeutic strategies. If the affinity-enhancing mutations are known, one may try to choose “cold-spots” for this mutation as targets for developing a possible therapeutic strategy instead of “hot-spots”. A comparison of hundreds of SARS-CoV-2 genome sequences suggests that the rate of non-synonymous mutations in SARS-CoV-2 is less than that of synonymous ones indicating that the active SARS-CoV-2 is at present under the control of negative selection. This is because presently, the virus is not under significant immune or drug pressure. If these pressures mount, the positive selection of non-synonymous mutations will accumulate much rapidly. This will cause a change in the viral phenotype allowing it to evade the host immune system and also develop drug resistance [33]. Thus, the weakening of the RBD-ACE2 complex interaction by modulating affinity-enhancing/affinity-weakening mutations represents a possible therapeutic strategy.

The rapid increase in the atomic-level understanding of molecular determinants desirable for the development of specific therapeutics has led to significant advances in the computational design of conformationally stable epitope-scaffolds and inhibitors. In recent years high-throughput computation and mutational mapping have been used to optimize the binding interface and develop several peptide-based inhibitors and conformationally stable epitope-scaffolds against several enveloped and non-enveloped viruses [34, 35]. These computationally designed, conformationally stable epitope-scaffolds were able to elicit superior epitope-specific responses compared to the wild-type viral protein [35–39]. We suggest that such protein-design based rational approaches may also be explored in designing next-generation precision vaccines and drugs against SARS-CoV-2 [40, 41].

## 5. CONCLUSIONS

The natural evolution of the SARS-CoV strains might have occurred upon selection over a long period of time, through the repeated transmission of viruses from animals to humans and vice-versa, resulting in mutations that favored SARS-CoV transmission and pathogenicity. Understanding of integrated system biology of biomolecular interaction networks during viral infection in the context of sequence variations in the viral membrane protein responsible for cell entry provides several crucial information, such as, i) understanding the clinical pathogenesis and progression of the virus, ii) regions/residues that may weaken the RBD/ACE2 interaction, iii) development of specific fusion-inhibiting therapeutics, such as small molecule peptides and non-peptidic inhibitors, and iv) design and development of monoclonal antibodies targeting RBD that can serve as immunotherapeutic or passive immunization agents. In conclusion, our work offers a cohesive and comprehensive picture of the mutational landscape of SARS-CoV-2-RBD, providing information on both molecular mechanisms of infection and therapeutic routes useful for treating the COVID-19 outbreak.

## ACKNOWLEDGMENTS

The authors acknowledge RIKEN ACCC for the Hokusai supercomputing resources.

## COMPETING INTERESTS

The authors have declared no competing interest.

## AUTHORS’ CONTRIBUTIONS

AKP carried out the experiments and data generation. AKP and TT conceived the study and participated in its design and coordination. AKP, PK, KYJZ, and TT analyzed the data and drafted the manuscript. All authors read and approved the final manuscript.

