## Supplementary Tables for "High Throughput Designing and Mutational Mapping of RBD-ACE2 Interface Guide Non-Conventional Therapeutic Strategies for COVID-19"

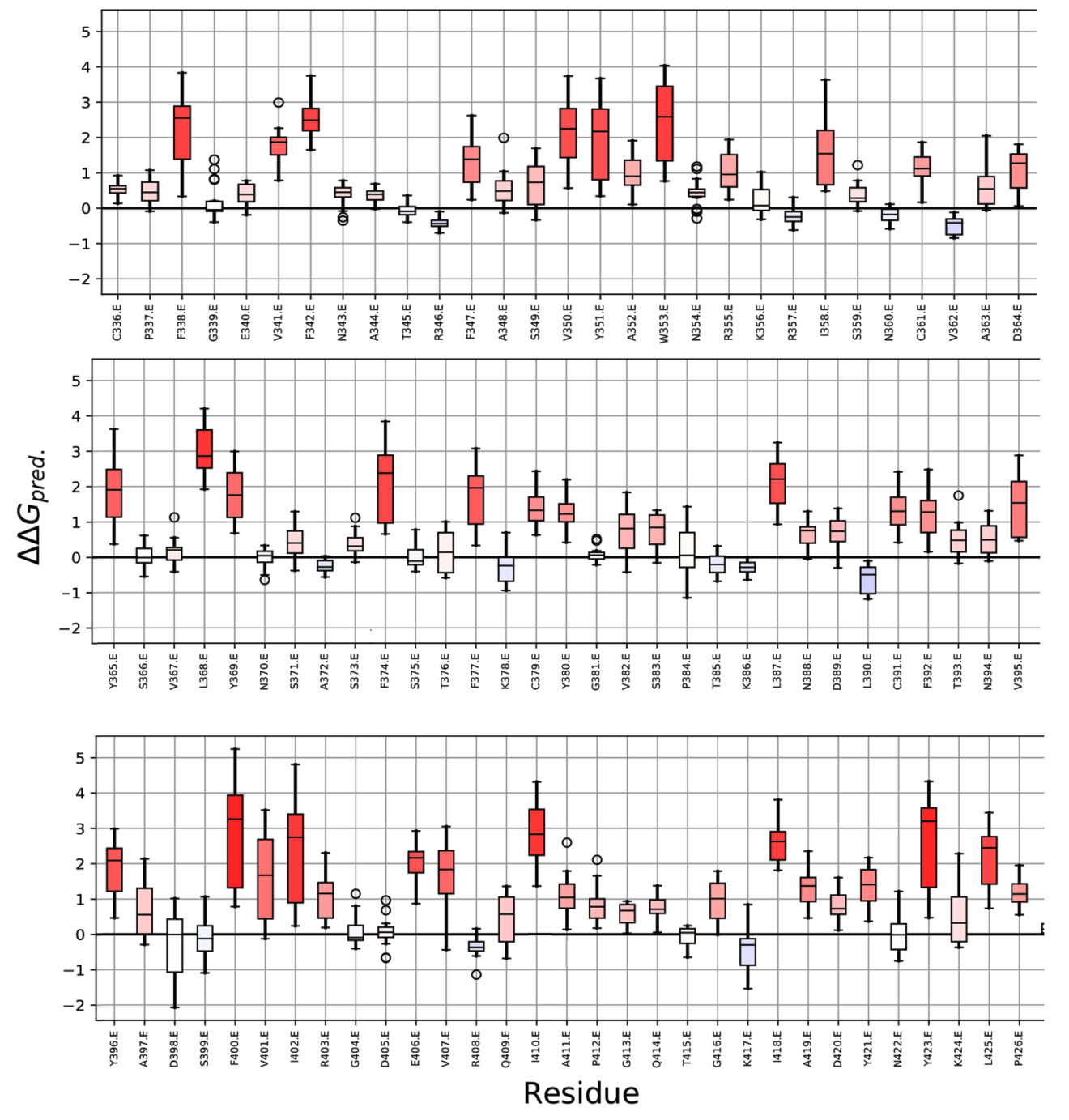

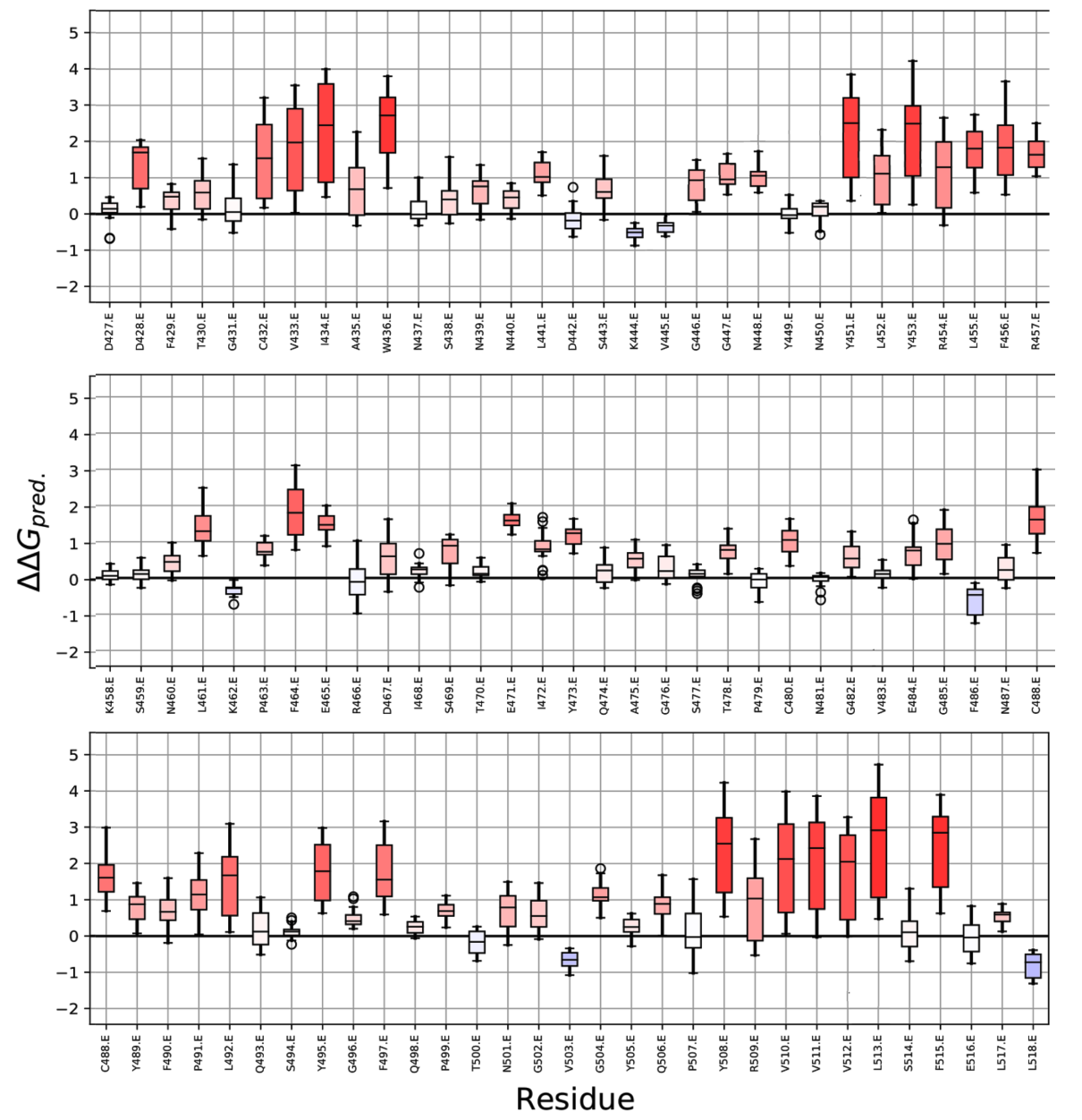

**Supplementary Figure S1.** Mutation sensitivity profile of each of the RBD residues is shown, with ΔΔG_pred_>0 or ΔΔG_pred_<0 demonstrating the impact of each mutation at individual positions.

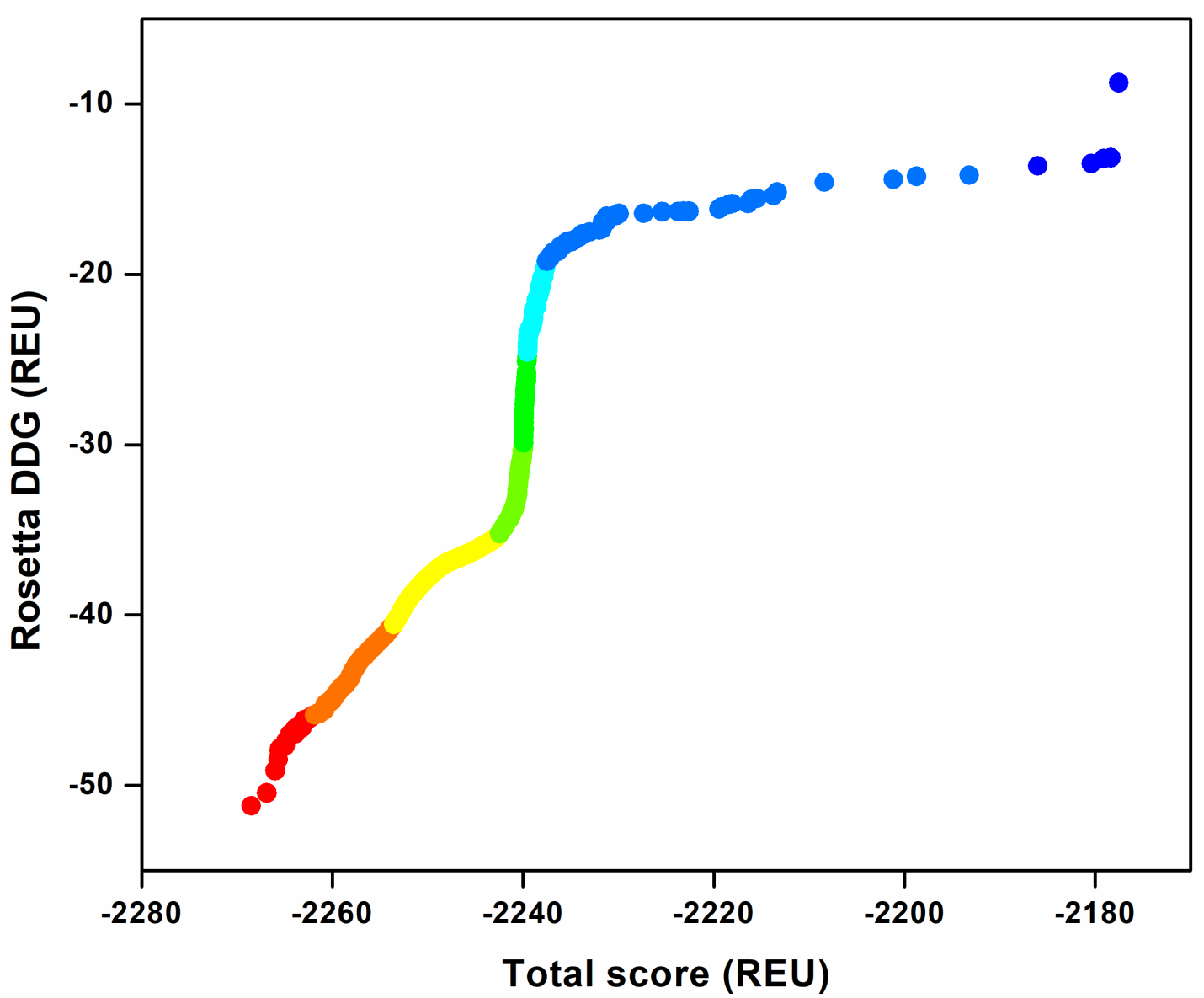

**Supplementary Figure S2.** Rosetta total score (REU) versus Rosetta DDG (REU) for the computed designs showing the convergence of design calculations. The Rosetta total score and Rosetta DDG show a well converged trajectory from lower to higher scores and binding affinities, respectively.

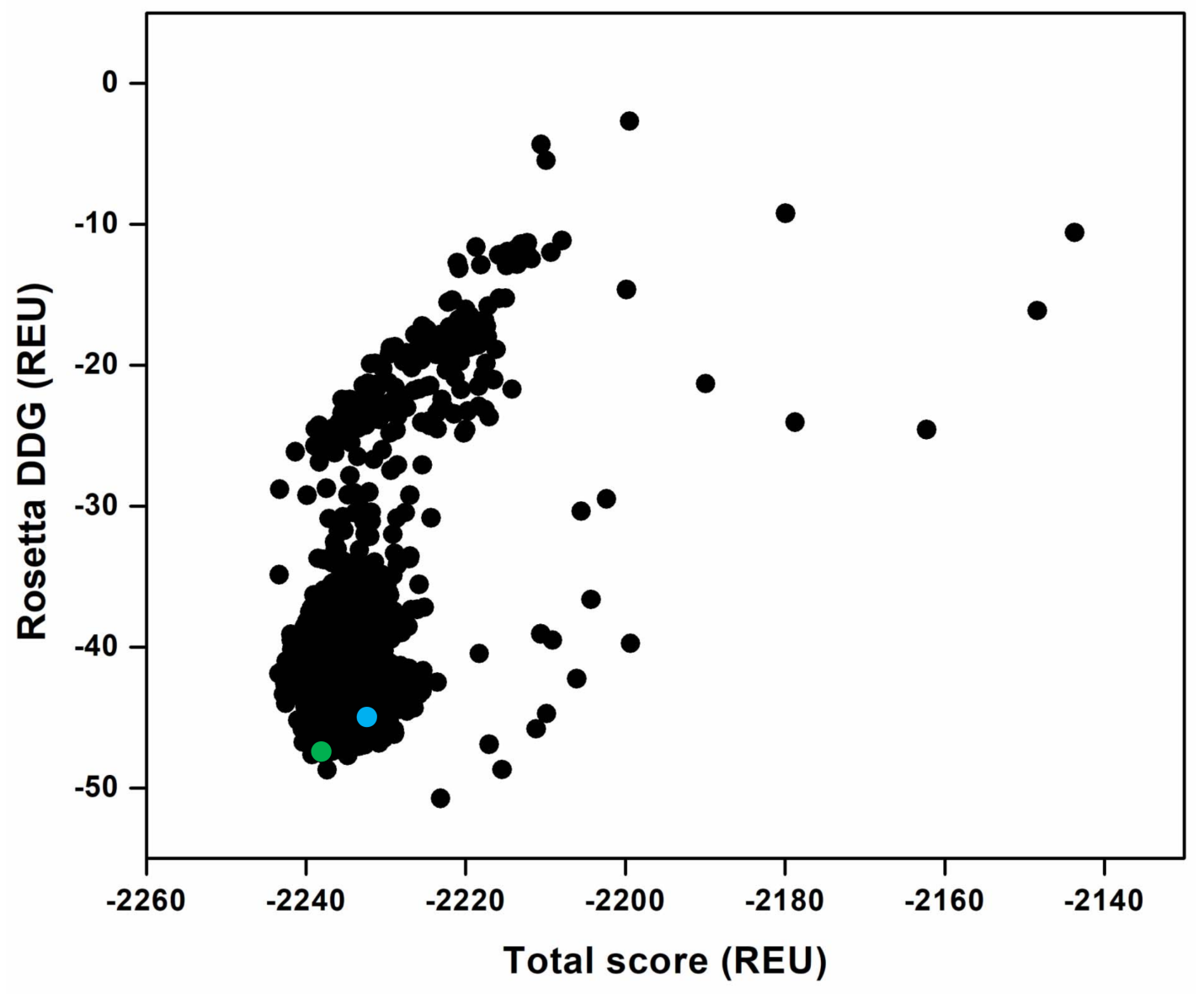

**Supplementary Figure S3.** Rosetta total score (REU) versus Rosetta DDG (REU) for the computed designs where only G467 and V483 are designed. The G467S and V483A designs are among the top-ranked designs shown in green and blue colors, respectively.

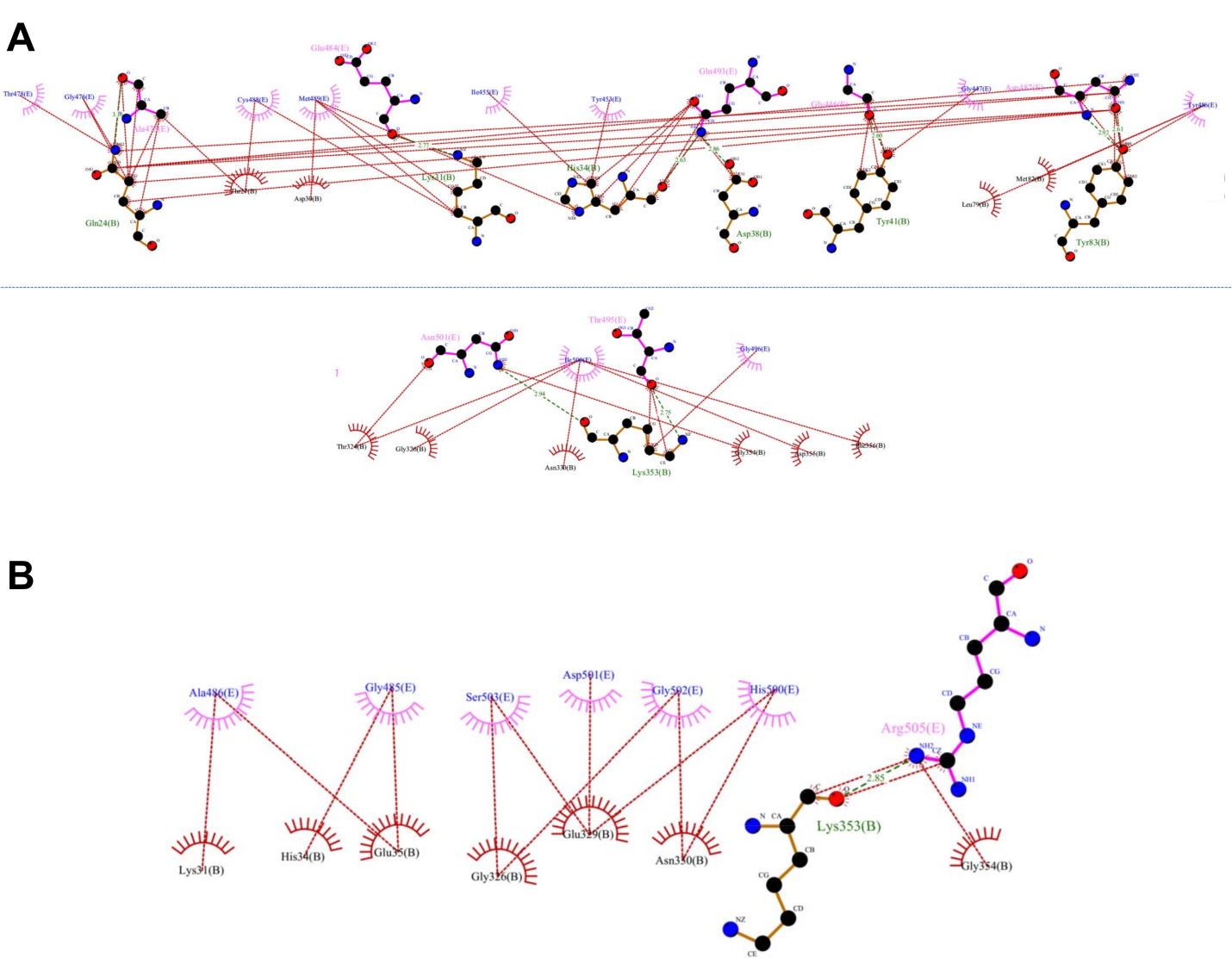

**Supplementary Figure S4.** The residue-level interactions between ACE2 and RBD for the (A) affinity-enhancing Design-1 and (B) affinity-weakening Design-1 obtained by the DIMPLOT program of LigPlot+v1.4.5 are shown indicating the higher number of interactions in (A). Because of the extended interface, the interactions in panel (A) are splitted for better clarity, and the horizontal dashed line represents the continuity of the interactions.
